# *Cis*-regulatory changes in locomotor genes are associated with the evolution of burrowing behavior

**DOI:** 10.1101/2021.09.27.462036

**Authors:** Caroline K. Hu, Ryan A. York, Hillery C. Metz, Nicole L. Bedford, Hunter B. Fraser, Hopi E. Hoekstra

## Abstract

How evolution modifies complex, innate behaviors is largely unknown. Divergence in many morphological traits has been linked, at least in part, to *cis*-regulatory changes in gene expression, a pattern also observed in some behaviors of recently diverged populations. Given this, we compared the gene expression in the brains of two interfertile sister species of *Peromyscus* mice, including allele-specific expression (ASE) of their F_1_ hybrids, that show large and heritable differences in burrowing behavior. Because *cis*-regulation may contribute to constitutive as well as activity-dependent gene expression, we also captured a molecular signature of burrowing circuit divergence by quantifying gene expression in mice shortly after burrowing. We found that several thousand genes were differentially expressed between the two sister species regardless of behavioral context, with several thousand more showing behavior-dependent differences. Allele-specific expression in F_1_ hybrids showed a similar pattern, suggesting that much of the differential expression is driven by *cis*-regulatory divergence. Genes related to locomotor coordination showed the strongest signals of lineage-specific selection on burrowing-induced *cis*-regulatory changes. By comparing these candidate genes to independent quantitative trait locus (QTL) mapping data, we found that the closest QTL markers to these candidate genes are associated with variation in burrow shape, demonstrating an enrichment for candidate locomotor genes near segregating causal loci. Together, our results provide insight into how *cis*-regulated gene expression can depend on behavioral context as well as how this dynamic regulatory divergence between species can be integrated with forward genetics to enrich our understanding of the genetic basis of behavioral evolution.

## Introduction

Animals exhibit a diverse array of innate behaviors, the genetic basis of which remains poorly understood. Comparative and evolutionary developmental studies of morphology have linked numerous traits to differences in the *cis-*regulatory control of gene expression (Wittkopp et al., 2004). Given these patterns, it is likely that at least some components of behavioral evolution are associated with *cis*-regulatory changes (e.g. Wang et al., 2019; York et al., 2018). Technical advances are enabling behavioral and transcriptomic studies across an increasingly broad swath of animal species and clades (Jourjine and Hoekstra, 2021). Here we dissect gene-regulatory contributions to variation in an innate, natural behavior – burrowing – among deer mice (genus *Peromyscus*).

Deer mice are an emerging system for investigating the mechanisms of behavioral evolution (Bedford and Hoekstra 2015). Two sister species, *P. maniculatus* and *P. polionotus*, differ in their species-typical burrow size and shape. *P. maniculatus* dig short burrows (<10 cm), consisting of an entrance tunnel and a nest chamber while *P. polionotus* construct longer (>35 cm) burrows which, in addition to an entrance tunnel and nest chamber, also include an upward sloping “escape tunnel” (Figure 1A) (Dawson et al., 1988; Weber and Hoekstra, 2009). One explanation for this behavioral difference is that long burrows buffer against environmental fluctuations in the open habitats of *P. polionotus* (Bedford et al., 2021). Morphological comparisons between these two species have not found evidence for digging-related specializations (e.g., forepaw enlargement as seen in moles), suggesting burrow differences are largely driven by behavioral mechanisms (Hu and Hoekstra, 2016). Indeed, cross-fostering experiments further suggest that this behavioral variation has a strong genetic component (Metz et al., 2017).

**Figure 1.**
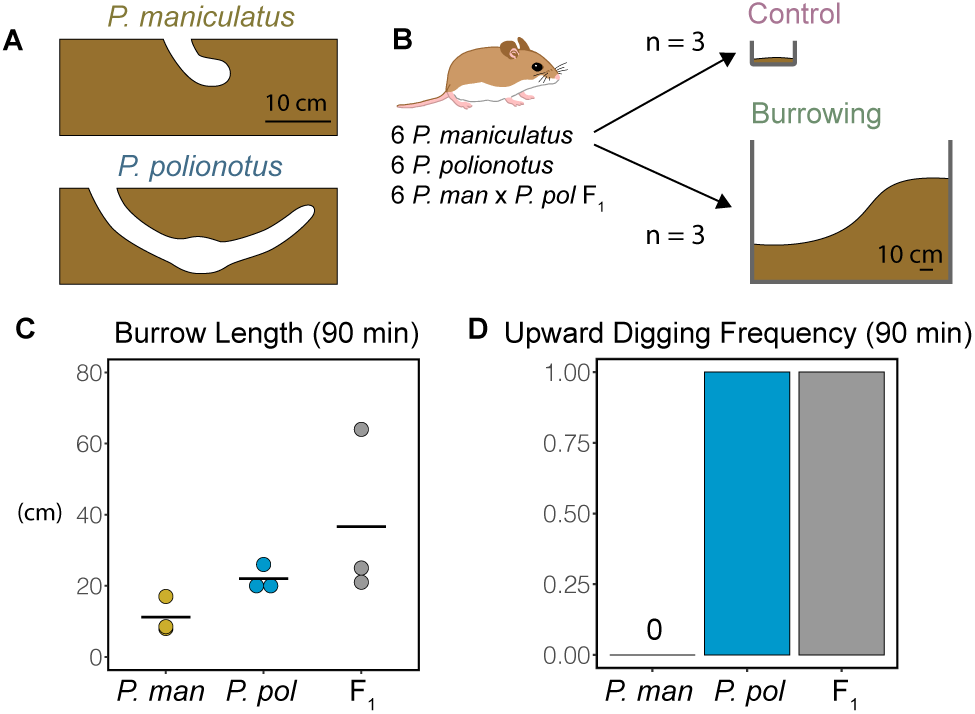
Burrowing behavior experimental design and phenotypes. **(A)** Species-typical burrow shapes of *P. maniculatus* and *P. polionotus*. **(B)** Mice were split into control and burrowing cohorts and exposed to the corresponding enclosure for 90 minutes prior to dissection. **(C)** Burrow lengths dug during 90-min test period by burrowing group. **(D)** Frequency of upward digging observed during 90-min test period by burrowing group.

A forward-genetic screen has shown that the *P. polionotus* burrow is dominantly inherited, and that genomic regions affecting divergence in burrow size and shape can be mapped to distinct loci (size: 3 loci, shape: 1 locus; (Weber et al., 2013)). However, identifying causal genes and pathways from such QTL analyses remains a major challenge. Furthermore, although a widespread behavior, very little is known about the genetic basis of burrowing, offering no obvious candidate genes.

To address this, we developed an integrative approach for identifying burrowing-associated genes and pathways via comparative transcriptomics. Although comparing gene expression across species can be informative, it is often affected by confounding factors such as differences in cell-type abundances, developmental timing, and the unavoidable variability that accumulates during an organism’s lifetime. Moreover, in such comparisons *cis*- and *trans*-acting regulatory divergence cannot be disentangled. However, measuring ASE in F_1_ hybrids circumvents these issues: it controls for these confounders because the alleles being compared are present in the same individuals in the same cells and only reflects *cis*-regulatory divergence (since *trans*-acting effects are expected to affect both alleles equally).

By analyzing neural gene expression in burrowing F_1_ hybrids of *P. maniculatus* (short burrow) and *P. polionotus* (long burrow), we identified extensive *cis*-regulatory differences related to behavioral state. Intersecting these results with QTL data connected a discrete subset of locomotor-related genes displaying species-specific expression with burrowing loci, implicating their involvement in the evolution of this complex behavior.

## Results

We first introduced all experimental mice (n = 18; 6 mice for each of three “genotypes”: *P. maniculatus, P. polionotus*, F_1_ hybrids) to a large sand-filled enclosure overnight and confirmed they dug full length genotype-typical burrows (Figure S1). We observed a pattern of *P. polionotus-*dominant burrow trait inheritance in F_1_ hybrids that is consistent with previous studies (Dawson et al., 1988; Weber et al., 2013).

To measure burrowing-induced gene transcription, we then exposed a “test” cohort of mice (n = 9; 3 mice per genotype) to large sand enclosures for 90mins. This timeframe captures the rise in primary and secondary response in gene transcription to a stimulus (i.e. genes that do not need *de novo* translation for transcription and the following wave) (Tullai et al., 2007). A second “control” cohort of animals (n = 9; 3 mice per genotype) was exposed to a thin layer of sand in a new housing cage to account for handling and sensory stimuli from sand (Figure 1B). In this 90-minute period, we found all nine “test” mice had burrowed substantially, although none to the full burrow length achieved in overnight trials (Figure 1C, Table S1). Importantly, all *P. polionotus* and F_1_ hybrids – but not *P. maniculatus* – had begun upwards digging that is characteristic of the escape tunnel (Figure 1D). Thus, the mice included in this study, in both overnight and acute trials, behaved in a genotype-typical way.

To measure overall and burrowing-dependent expression divergence between *P. maniculatus* and *P. polionotus*, we used RNA-sequencing (RNA-seq) to generate whole brain transcriptomes from both burrowing test and control animals that had not burrowed. Because gene expression depends on cell type (Hrvatin et al., 2018) as well as the pattern of activity (Tyssowski et al., 2018), we anticipated neural activity specific to burrowing would affect gene expression. Furthermore, since neural substrates underlying burrowing have diverged between these two species, we expected to find species-specific differences in burrowing-induced gene expression. We first computed differential expression between the two species (Figure 2A; see Methods for all comparisons tested). We detected approximately 14,000 expressed genes (14,126 for burrowing mice and 14,393 for control animals; Transcripts per million [TPM] >1), and 13,273 genes were shared across all 12 samples. Of those, we found a total of 3,619 genes were differentially expressed between species in both the burrowing and control conditions, and an additional 1,962 and 670 genes to be differentially expressed in either only the burrowing or control conditions, respectively (Benjamini-Hochberg adjusted *P <* 0.05; Figure 2B). Principal component analysis of whole transcriptome data cleanly separated *P. maniculatus* and *P. polionotus* along PC1 (86.95% of variance explained; Figure 2C). These results demonstrate that, while the majority of differentially expressed genes were stable across treatments, a subset of genes show species-specific regulation in response to behavioral condition.

**Figure 2.**
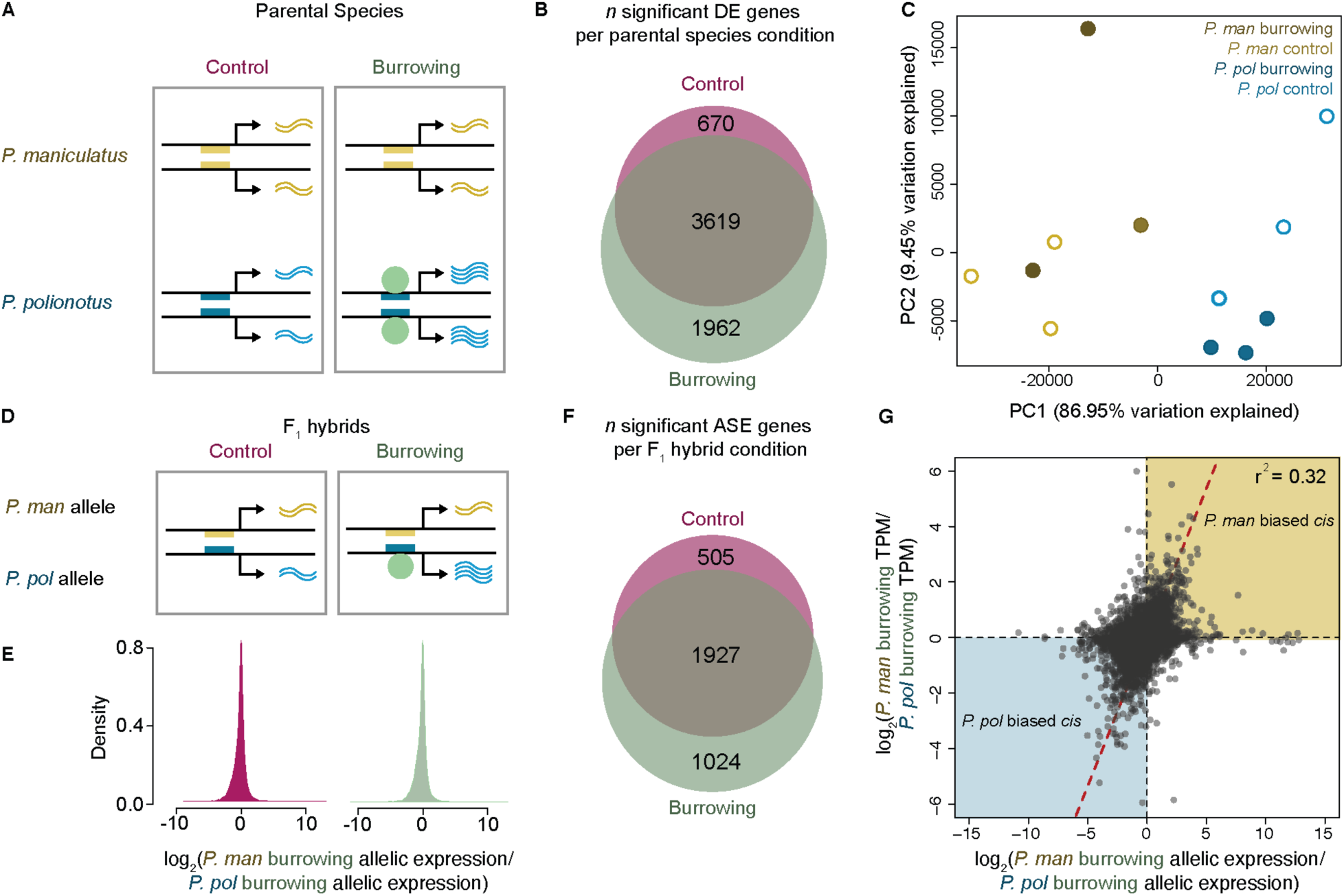
Gene expression divergence between *P. maniculatus* and *P. polionotus* alleles. **(A)** Schematic of potential mechanism underlying behavior-dependent differential expression between *P. maniculatus* and *P. polionotus*. Gene regulatory regions corresponding to the *P. maniculatus* (yellow rectangle) and *P. polionotus* (blue rectangle) alleles. Species-specific sequence changes to this region may alter binding of a burrowing-responsive transcription factor (green circle). **(B)** Venn diagram of the number of the genes with significant differential expression between control *P. maniculatus* and *P. polionotus* (pink) and burrowing *P. maniculatus* and *P. polionotus* (green). **(C)** Principal component analysis of burrowing and control *P. maniculatus* (yellow) and *P. polionotus* (blue) whole brain transcriptomes. **(D)** Schematic of potential mechanism underlying allele-specific expression (ASE). **(E)** Distribution of allelic expression bias for all genes in control versus burrowing F_1_hybrids. **(F)** Venn diagram of the number of the genes with significant ASE between control F_1_ hybrids (pink) and burrowing F_1_ hybrids (green). **(G)** Scatterplot comparing the distribution of parental allele expression in burrowing F_1_ hybrids (x-axis) and gene expression in burrowing *P. maniculatus* and *P. polionotus* (y-axis). Identity line shown in red.

To investigate regulatory divergence in the neural transcriptomes of *P. maniculatus* and *P. polionotus*, we generated whole brain RNA-seq from *P. maniculatus* x *P. polionotus* F_1_ hybrids that were exposed to burrowing or control conditions. Measuring gene expression in F_1_ hybrids controls for differences in environment and cell-type abundances, which may contribute to expression differences in parental species, allowing for more direct comparison of parental alleles as well as the detection of allele-specific expression (ASE) indicative of *cis-*regulatory divergence (Fraser et al., 2011; Wittkopp et al., 2004). By extending analyses of ASE to multiple conditions, one can measure differential allele-specific expression (diffASE), which can identify context-dependent regulatory differences (York et al., 2018). In the case of behavior, such context-dependent regulatory divergence may reflect neural-activity-responsive genes that are controlled by regulatory elements containing sequence differences between the parental species (Figure 2D).

Our analyses of F_1_ hybrid transcriptomes identified ASE differences in genes from both burrowing and control animals. Overall, the distribution of ASE was slightly biased toward *P. polionotus* alleles (Figure 2E), which was not attributable to mapping bias because we used the *P. maniculatus* genome as our reference. Instead, this slight bias may reflect a global imbalance favoring paternal alleles as described in *Mus musculus* (Crowley et al., 2015), or could reflect a species-specific expression bias. Overall, we detected a total of 3,456 genes with significant ASE (Binomial test; Bonferroni adjusted *q* < 0.05). Like differential expression in the parental species, most ASE genes (n = 1,927) were shared between the two conditions; nonetheless a substantial portion was found only in burrowing animals (n = 1,024) and to a lesser extent only in control animals (n = 505) (Figure 2F). A comparison of the distribution of F_1_ hybrid allelic ratios during burrowing (log_2_[*P. maniculatus* allele*/ P. polionotus* allele]) to the ratios of parental species burrowing expression (log_2_[*P. maniculatus* TPM*/ P. polionotus* TPM]) indicated the presence of both *cis-* and *trans-*regulatory differences (Figure 2G). The same comparison between control F_1_ hybrid samples and control parental species samples yielded similar results (Figure S2). Accordingly, the overall relationship between the two distributions (*r*^*2*^ *=* 0.32) was similar to that observed in other studies of interspecific hybrids produced by crossing diverged lineages (e.g. Goncalves et al., 2012; Mack et al., 2016; McManus et al., 2010). Taken together, these observations suggest that detectable regulatory differences in *Peromyscus* brain gene expression arise from both *cis-* and *trans-*acting divergence, and that capturing the brain in different behavioral states can unmask further regulatory differences. Our data suggest that *cis*-regulation in the brain can be highly context-specific (York et al., 2018), in contrast to other tissues or species where *cis*-regulation is robust to environmental changes (Verta and Jones, 2019).

Given these observations, we next formally tested for the presence of context-dependent *cis*-regulation. First, we identified genes with differential ASE (diffASE), for example, those that showed allele-specific induction or suppression between burrowing and control F_1_ hybrids (Figure 3A). Comparing mean levels of ASE between burrowing and control hybrids identified a number of outlier genes (Figure 3B) despite the fact that, overall, ASE values were relatively consistent between the two conditions (*r*^*2*^ = 0.94). To test for the presence of diffASE, we first filtered our data to include only genes with evidence of expression across all six individuals (11,928 genes) and then used a Fisher’s exact test to compare allelic counts in a 2×2 contingency table (*P. polionotus* burrowing allele, *P. polionotus* control allele; *P. maniculatus* burrowing allele, *P. maniculatus* control allele) for pairs of burrowing and control samples (see Methods). The resulting p*-*values were combined using Fisher’s method and adjusted via Bonferroni correction. After doing so, we detected 2,844 genes with significant combined diffASE. We further narrowed this list to those genes possessing significant diffASE across all three burrowing and control pairs, resulting in a conservative and high confidence list of genes displaying diffASE (n = 177 genes; Figure 3B). While the majority of these genes retained their direction of bias between burrowing and controls, a subset did switch from *P. maniculatus*-biased in control samples to *P. polionotus*-biased in burrowing samples (n = 24/177), while no genes switched in the opposite direction, a statistically significant difference (Fishers’ exact test: *P* = 5.17×10^−8^). Together, these data identify genes that change expression level in mice when they burrow and show that we see more upregulation of alleles from the long-burrowing *P. polionotus* than from the short-burrowing *P. maniculatus*.

**Figure 3.**
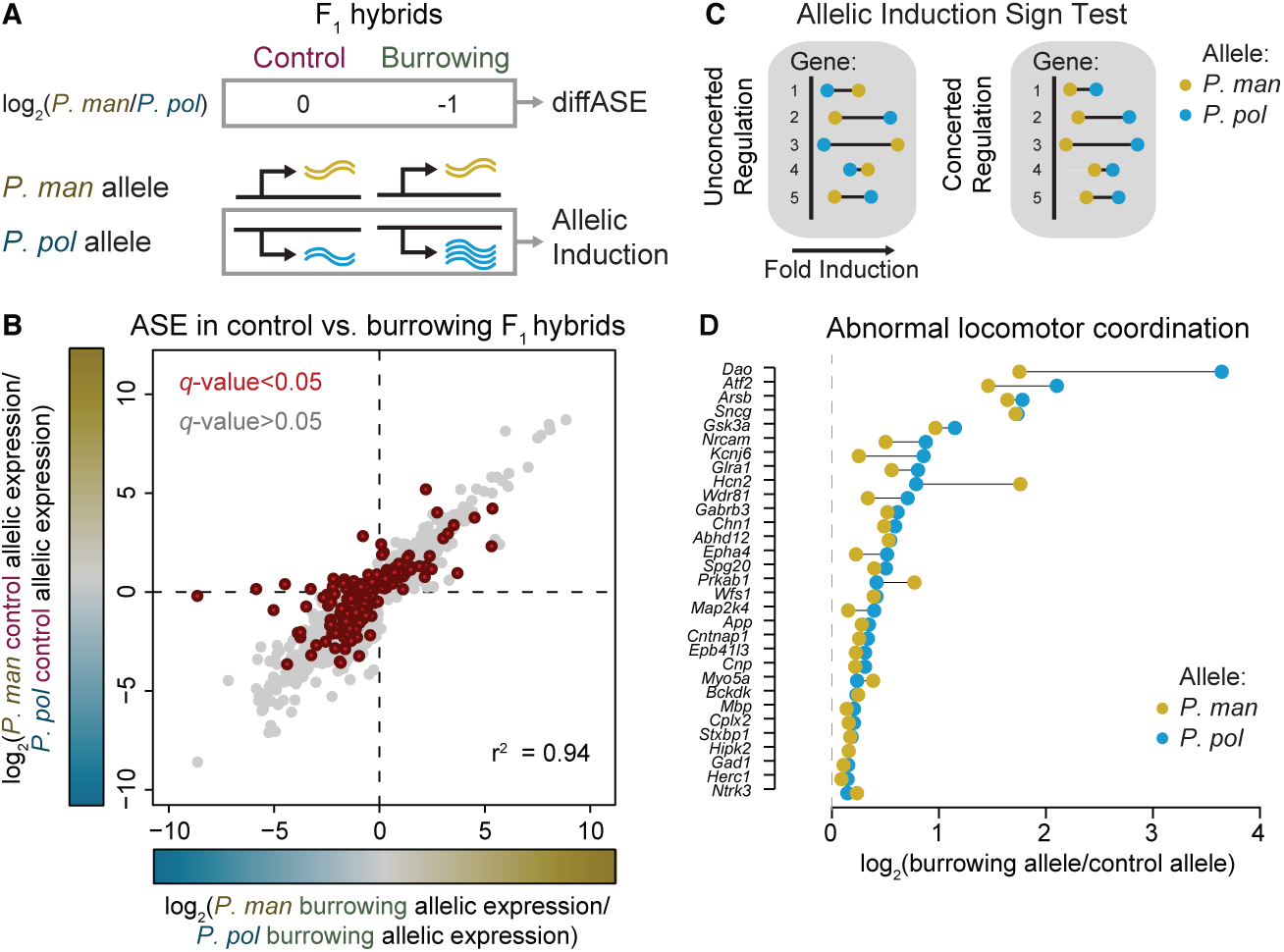
Analyses of context-dependent allele-specific expression. **(A)** Schematic showing comparison between F_1_ behavioral groups to calculate differential allele-specific expression (diffASE) (top) and allelic induction (bottom). **(B)** Scatterplot comparing the distribution of allelic ratios between burrowing and control conditions. Points (representing individual genes) are colored by binned diffASE *p*-values obtained using a Fisher’s exact test on *P. maniculatus* and *P. polionotus* allele counts in the two conditions. **(C)** Schematic outlining the logic of the sign test. Here we consider a functional set composed of 5 genes with corresponding values for allelic induction (the magnitude of which is represented by distance from the y-axis). Under unconcerted regulatory evolution, we expect the parental allele with greater induction to be relatively evenly shared between the two alleles (left). In the case of concerted regulatory evolution, we expect a strong preference in allelic induction toward one species or the other (right). **(D)** Scatterplot highlighting the distribution of allelic induction for diffASE genes in the locomotor coordination functional category.

We next asked whether genes exhibiting burrowing-dependent diffASE could implicate functional mechanisms underlying burrowing evolution. Genes involved in a specific pathway or biological process can exhibit concerted gene expression (i.e. the genes will be transcriptionally up-regulated or down-regulated together) (Fraser et al., 2011; York et al., 2018). Because burrowing predominantly was associated with an increase in gene-expression level, we focused on whether a diffASE gene exhibited greater allelic induction in one parental allele than the other (Figure 3A). If a specific pathway or biological process has undergone selection, then we expect an enrichment of alleles with the same direction of allelic induction. To test this hypothesis, we employed the sign test, a framework for detecting lineage-specific selection on the regulation of functionally related groups of genes (as defined by the Mammal Phenotype Ontology (Smith and Eppig, 2009)). We tested seven gene sets for whether member genes displayed consistent biases in allelic induction toward one parental species or the other (Fraser et al., 2011; York et al., 2018) (Figure 3C; see Methods). We found that significantly more *P. polionotus* alleles involved in abnormal locomotor coordination (MP:0001392) were upregulated during burrowing compared to the *P. maniculatus* allele (n = 26/31 alleles; Bonferroni corrected Fisher’s exact test *P <* 0.001; Figure 3D). The second strongest signal of concerted regulation also involved a category related to locomotor defects: abnormal gait (MP:0001406; Bonferroni corrected Fisher’s exact test *P* < 0.06), although not statistically significant. Thus, we find evidence that the dynamic, behavior-dependent *cis*-regulation of genes involved in locomotion was subject to lineage-specific selection between *P. polionotus* and *P. maniculatus*.

Having identified locomotion-related gene sets with behavior-dependent *cis*-regulation, we next explored whether these genes are strong candidates for harboring causal mutations underlying species differences in burrowing behavior or, alternatively, simply downstream genes responding to burrowing. To test this, we exploited a published QTL study for burrow traits in a large backcross between *P. polionotus* and *P. maniculatus*, which identified three genomic regions associated with burrow architecture (Weber et al., 2013). However, additional causal genetic variants likely contribute to differences in burrowing behavior, but have effect sizes too small to reach genome-wide significance as QTL, should nevertheless be enriched for some degree of genetic association with burrow traits. Therefore, we determined if the genetic markers nearest our candidate genes affecting locomotor coordination are associated with traits such as burrow size, shape, and escape tunnel presence/size (Figure 4A).

**Figure 4.**
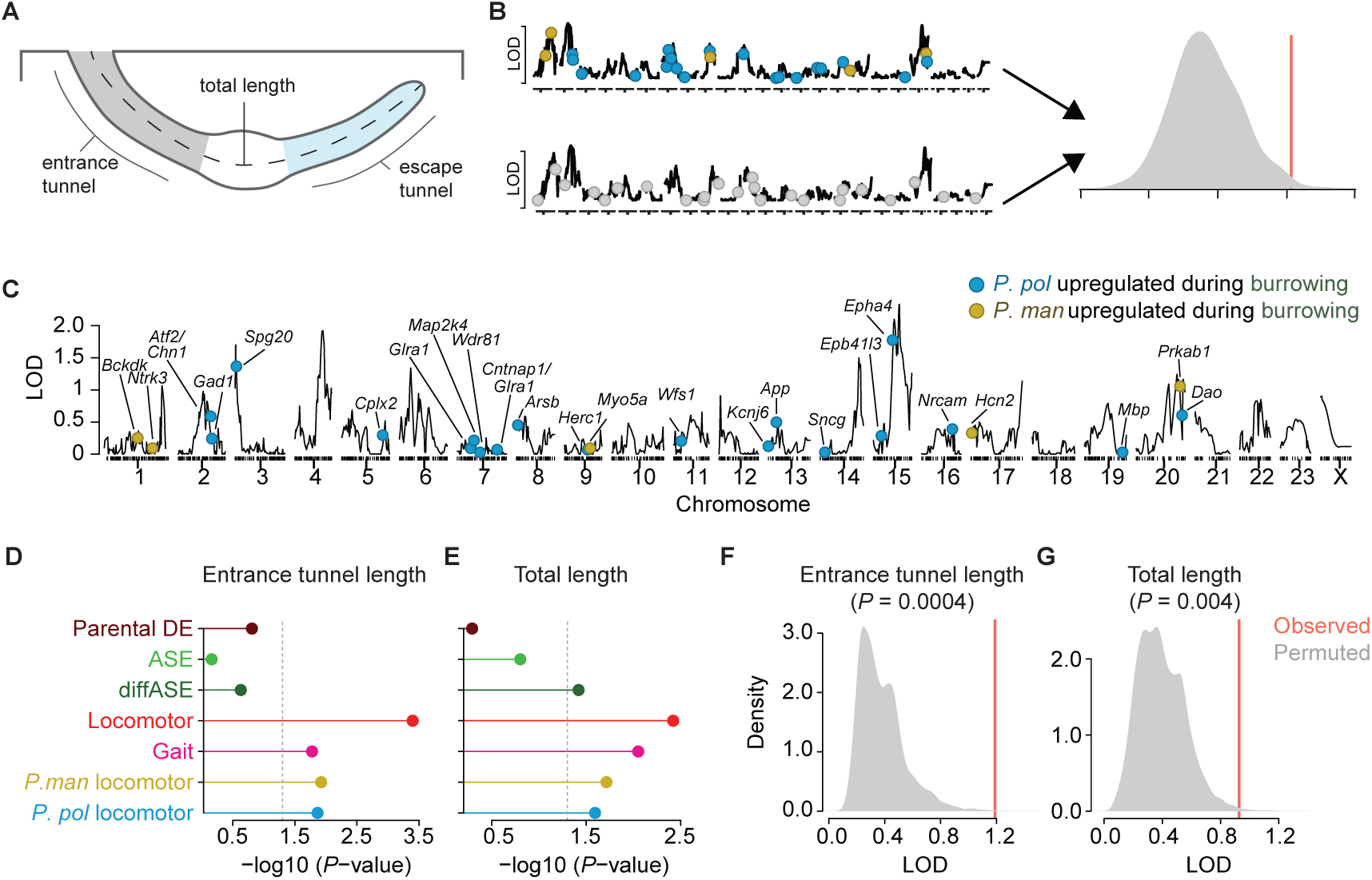
Intersection of QTL and diffASE. **(A**) Diagram of a burrow and measured traits. **(B)** Schematic of the procedure for permutation testing of QTL/diffASE intersection. Model fits for gene markers from the list of interest and 10,000 random sets of the same size are computed (represented by LOD plots). The observed fit (red line) is then compared to the 10,000 random sets (grey distribution) to calculate a *P-*value. **(C)** Plot of genome-wide LOD scores for entrance-tunnel length (taken from Weber et al. 2013). Overlaid are the locations of the closest markers to genes in the abnormal locomotor coordination set colored by species. **(D)** and **(E)** Scatterplots of results for selected measures from the permutation test. For each measure -log10(*p*-values) are plotted for the following subsets: 200 most differentially expressed genes between *P. maniculatus* and *P. polionotus* while burrowing (brown; ‘Parental DE’), genes with significant ASE (light green; ‘ASE’), genes with significant diffASE (dark green; ‘diffASE’), diffASE genes associated with locomotor function (red; ‘Locomotor’), diffASE genes associated with gait (magenta; ‘Gait’), *P. maniculatus*-biased locomotor diffASE genes only (yellow; ‘*P. man* motor’), and *P. polionotus*-biased locomotor diffASE genes only (blue; ‘*P. pol* motor’). **(F)** and **(G)** histograms of permutation test results for diffASE genes associated with locomotor function for the same traits shown in **(D)** and **(E)**, respectively.

In addition, we sought to compare the QTL associations of the differentially induced locomotor-coordination genes with other forms of gene expression divergence that are more commonly analyzed (e.g. allele-specific expression in hybrids or differential expression between species). If causal genetic differences for a given trait are associated with differential allelic induction, then we expect that the LOD distributions for the candidate locomotor genes should be greater than the other sets. To test this, we assessed the LOD distributions of markers representing a range of gene-expression divergence types: genes associated with locomotor coordination (*P. maniculatus* [n = 5 genes] or *P. polionotus* [n = 26] biased genes only) and abnormal gait (n = 22), genes with significant diffASE (n = 177), genes with significant ASE (n = 200), and genes most differentially expressed between *P. maniculatus* and *P. polionotus* while burrowing (n = 200). While a high log odds (LOD) score near a gene certainly does not prove the involvement of that specific gene in the trait, observing a systematic bias towards high LOD scores for a group of genes strongly implies that group is enriched for genes affecting trait divergence.

When we compared the LOD scores of each burrow trait for markers closest to differentially induced locomotor-coordination genes to those of randomly permuted marker sets (Figure 4B and 4C), we found that the candidate markers had significantly higher LOD scores than the random sets for average entrance tunnel length (Figures 4D, 4F; *P* = 0.0004; 10,000 permutations, see Methods), maximum entrance tunnel length (*P =* 0.006), and average total length (Figures 4E, 4G; *P =* 0.004). These results suggest that this gene set is enriched for causal genetic effects relative to the genomic background (Table S2; with the caveat that other causal genes not identified by our RNA-seq analysis could also be nearby our selected markers; however this should only add noise, making our results conservative). Furthermore, locomotor-coordination genes tended to display stronger associations compared to all other gene sets across the traits tested (Figures 4D-E).

Given the association between the locomotor-coordination gene set and burrowing phenotypes, we next asked whether this association is affected by the direction of gene expression bias. For example, because *P. polionotus* burrow-shape traits appear largely dominant, we may expect animals with more *P. polionotus* genotypes at diffASE genes to have increased trait values. As described above, regression-based tests of the genotype-phenotype relationships demonstrated associations with diffASE genes involved in locomotor coordination and total burrow length, entrance tunnel length, and escape tunnel length (Figure 4D; Table S3). Within this group of genes, *P. polionotus*-biased genes, in particular, showed a stronger association with escape tunnel length than the other two traits. Escape tunnel is a trait for which previous QTL mapping identified only a single peak (with escape tunnel presence/absence treated as a binary trait) (Weber et al., 2013); however, our ASE approach implicated additional candidate genes residing in regions of moderate trait association in the genome. Long entrance tunnels and the presence of an escape tunnel are derived traits in *P. polionotus* (Weber and Hoekstra, 2009), and these results suggest these two traits are likely related to *P. polionotus*-specific changes to locomotor control.

## Discussion

Burrowing is a natural, complex, innate behavior – comprised of a series of coordinated head and limb movements – that consistently differs even among closely-related species of *Peromyscus* (Hu and Hoekstra, 2016). At the extreme, *P. polionotus* uniquely constructs long burrows, including an escape tunnel. QTL mapping of burrow traits revealed three genomic loci that harbor mutations for the derived burrow architecture observed in *P. polionotus* (Weber et al., 2013). Here, to move from large QTL regions to specific genes, we developed a complementary RNA-seq based strategy to identify patterns of gene expression, and ultimately candidate genes, that show lineage-specific changes in the *P. polionotus* brain. We find that genes associated with locomotor coordination have undergone *cis*-regulatory change such that, for the majority of locomotor candidate genes, the *P. polionotus* allele is induced while animals are burrowing. While we cannot conclude that these changes are directly linked to burrow evolution, since there are other ecological changes that may select for differences in locomotion, we do show that these candidate genes are enriched for association with burrow traits in a genetic cross, suggesting that they may influence the evolution of this “extended phenotype”. We should emphasize that, while this finding may seem intuitive, it would not have emerged by analyzing parental differential expression or “baseline” allele-specific expression alone. Instead, we were able to implicate candidate genes, even those which may have only a small behavioral effect, through our approach, using only a fraction of the animals that would be needed for a QTL-mapping population. Thus, by measuring differential (i.e., behavior-evoked) allele-specific expression, we can identify genes (and pathways) with dynamic *cis*-regulatory divergence that likely contribute to an ecologically important and complex behavior.

## STAR Methods

### Key Resources Table

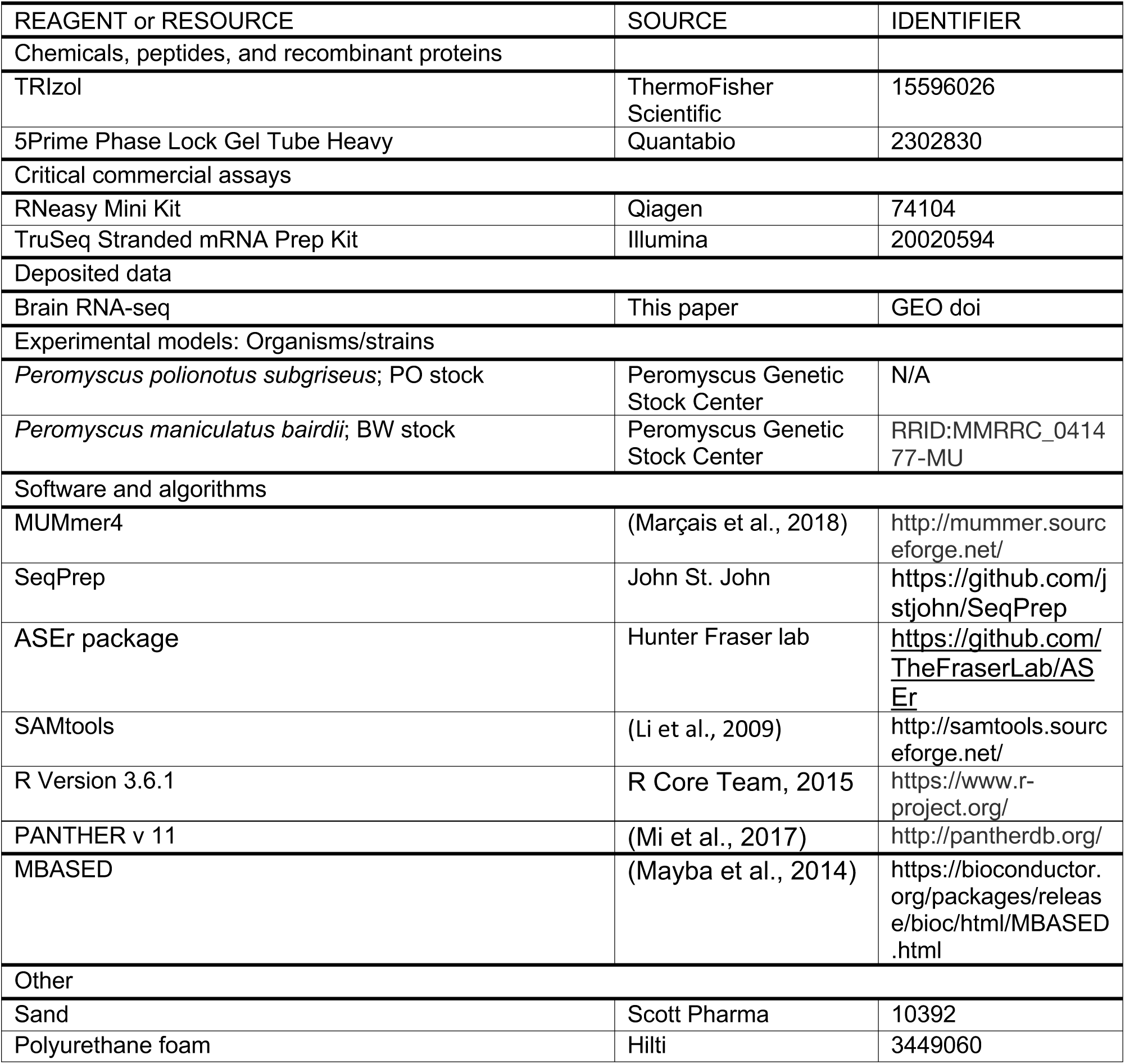

### Resource Availability

#### Lead contact

Further information and requests for resources and reagents should be directed to and will be fulfilled by the lead contact, Hopi E. Hoekstra.

### Materials availability

This study did not generate new unique reagents.

### Data and code availability

- RNA-seq data have been deposited at GEO and are publicly available as of date of publication. Accession numbers are listed in the key resources table.
- This paper does not report any original code.
- Any additional information required to reanalyze the data reported in this paper is available from the lead contact upon request.

### Experimental Model and Subject Details

#### Animals

We originally obtained *P. maniculatus bairdii* (BW) and *P. polionotus subgriseus* (PO) outbred strains from the *Peromyscus* Genetic Stock Center (University of South Carolina, Columbia, SC, USA), and then established breeding colonies at Harvard University. We generated F_1_ hybrids by crossing *P. maniculatus* females with *P. polionotus* males; the reciprocal cross direction suffers from hybrid inviability (Maddock and Chang, 1979). All experimental animals were males and housed in standard polysulfone cages (19.7 × 30.5 cm and 16.5 cm high; Allentown, New Jersey, USA) in same sex and genotype groups, until testing adults at approximately 50-80 days of age. Housing cages contained enrichment as previously described (Lewarch and Hoekstra, 2018). We maintained animals at 22 °C with a 16:8h light:dark cycle and provided them with standard rodent food and water *ad libitum*. All procedures were approved by the Harvard University Institutional Animal Care and Use Committee (protocol 27-09-3).

## Method details

### Burrow phenotyping

We measured burrowing behavior using large sand-filled (Scott Pharma, Marlborough, MA, USA) enclosures (1.2 × 1.5 × 1.1 m) set up as previously described (Weber et al., 2013). We first ran all experimental animals through two “pretest” trials in the enclosures to allow them to acclimate and to confirm individuals burrowed in a species- (or hybrid-) typical manner. Next, we randomly assigned animals to a test trial cohort, “burrowing” or “control”. Trials were separated by a two-day rest period. For pretest trials, we released animals into enclosures two hours before the dark (active) phase of their light cycle. We then retrieved the animal 18 h later and returned it to its home cage. For the test trial, we released animals in the “burrowing” cohort into the large sand enclosures for 90 min. By contrast, we released animals in the “control” cohort into a new housing cage (as described above) containing only 0.14 kg sand, but also for 90 min. At the conclusion of each trial in the large sand enclosure, we identified all excavations with sufficient overhangs to conceal the animal’s body, classified them as “burrows”, and made a cast of the burrow with polyurethane foam (Hilti Corporation, Schaan, Liechtenstein) (Metz et al., 2017). We marked each cast with a horizontal level line, which was used to measure burrow length and determine whether an animal performed upward digging (i.e., an escape tunnel).

### RNA-seq library preparation

To capture behavior-relevant gene expression, we focused on the brain. Therefore, at the conclusion of the test trial, we immediately euthanized animals using CO_2_ inhalation and rapidly dissected whole brains in chilled PBS, flash-froze the sample in liquid nitrogen, and stored it at -80 °C. Later, we homogenized tissues using a TissueLyser (Qiagen, Venlo, Netherlands) in Trizol (ThermoFisher Scientific, Waltham, MA, USA), and extracted total RNA using 5Prime Phase Lock Gel Tubes Heavy (Quantabio, Beverly, MA, USA), followed by clean-up with RNeasy columns (Qiagen, Venlo, Netherlands). We prepared RNA-seq libraries with a TruSeq Stranded mRNA Library Prep Kit, following manufacturer’s instructions (Illumina, San Diego, CA, USA), and assessed library quality prior to sequencing using a TapeStation 2200 (Agilent Technologies, Santa Clara, CA, USA). We performed paired-end sequencing (2 × 125 bp) on the Illumina HiSeq platform (San Diego, CA, USA).

### Parental species RNA-seq

To compare gene expression both across treatments and species, we first removed low-quality and adaptor sequences using SeqPrep (https://github.com/jstjohn/SeqPrep). We then aligned these filtered reads to the *P. maniculatus* transcriptome (Pman_1.0_mRNA.fa) and quantified estimated expression using kallisto (Bray et al., 2016). We summed read counts across transcripts to obtain gene-level expression values and selected genes with expression > 1 count in at least one species. To compute scale normalized expression values across samples, we used TMM normalization. We then computed differential expression with the limma R package using voom transformation and linear models alternately using species or condition (burrowing/control) as factors (Ritchie et al., 2015). We used Benjamini-Hochberg correction (Benjamini and Hochberg, 1995) on the resulting *p*-values and considered genes with FDR < 5% as significant. Gene set enrichments were conducted using PANTHER v 11 (Mi et al., 2017).

### SNP calling for allele-specific analyses

To assign alleles to one of the two species in our F_1_ hybrid RNAseq data, we first obtained the *P. maniculatus* (Pman_1.0, GenBank accession number GCA_000500345.1) and *P. polionotus* (Ppol_1.3, GenBank accession number GCA_003704135.1) genomes. We called heterozygous SNPs between these two species’ genomes using the MUMmer toolkit (Marçais et al., 2018). First, for memory efficiency, we split the reference genome (*P. polionotus*) into ten approximately equally sized segments, and then aligned the *P. maniculatus* genome to each segment using the nucmer function. All called SNPs were obtained with the function show-snps (options: -I (remove INDELs); -r (sort by reference sequence); -l (include sequence length information)). Next, we combined the resulting 10 files containing SNP calls and filtered out ambiguous SNP calls. We used this “less conservative” list to mask the reference genome during alignment for downstream allele-specific expression analyses (see below). Again, using the function show-snps, we produced a second, “more conservative” list, this time obtaining all called SNPs and INDELs between the two species (options: -c [uniquely aligned sequences only]; -r [sort by reference sequence]; -l [include sequence length information]). To control for SNPs arising from genomic rearrangements between *P. maniculatus* and *P. polionotus*, we removed SNPs within one read length of an INDEL. We then filtered out ambiguous calls and selected only SNPs occurring within annotated coding genes, resulting in a list of inter-specific heterozygous SNPs for use in conservative allele-specific expression quantification downstream.

### Allele-specific expression quantification

To quantify differences in allele-specific expression (ASE) levels in F1 hybrids, we first removed low-quality and adaptor sequences from all F_1_ hybrid RNA-seq samples using SeqPrep (https://github.com/jstjohn/SeqPrep), as with the parental species samples (described above). We then masked the *P. maniculatus* genome at the sites called in the “less conservative” list above using the perl script MaskReferencefromBED (https://github.com/TheFraserLab/ASEr) to avoid reference genome mapping bias. Reads from all F_1_ hybrid samples were aligned to this masked genome using STAR v 2.4.2a (Dobin et al., 2013) in 2-pass mode. We removed duplicate reads from the resulting .bam files and sorted by mate pair using SAMtools (Li et al., 2009). We determined ASE at the read-level using the script GetGeneASEbyReads in the ASEr package (https://github.com/TheFraserLab/ASEr). To ensure confidence in the species of origin for each quantified SNP, we considered ASE at only sites provided in the “more conservative” list of heterozygous SNPs identified in the above section.

### Allele-specific expression analyses

To test for allele-specific expression differences, we first filtered ASE calls from GetGeneASEbyReads requiring at least 1 count per allele from all genes per sample. This was motivated by the fact that the present of 0 counts for an allele may arise from a preponderance of false negatives due to our use of a conservative and small, but high confidence, list of SNPs for the initial quantification of ASE. This choice reflects a tradeoff in the ability to find true monoallelic expression and the capacity to confidently assign reads to a species of origin, the latter of which we opted to prioritize given the broader evolutionary focus of this study. In addition to this initial filtering, we removed genes known to be imprinted in *Mus musculus* (Perez et al., 2015) (Table S4). After filtering we computed an ASE ratio for each gene by calculating the log_2_ ratio of *P. maniculatus* allelic counts compared to *P. polionotus* allelic counts. Positive values correspond to a *P. maniculatus* allelic bias while negative values reflect a *P. polionotus* bias.

To test for significant biases in ASE per sample, we used a two-tailed binomial test of the *P. maniculatus* and *P. polionotus* allele counts per gene and adjusted the resulting *p*-values within each sample using Bonferroni correction. We considered genes to show significant ASE in a given condition (control/burrowing) if they had adjusted *p-*values <0.05 across all three condition replicates and biased in the same direction.

### Calculating and analyzing differential allele-specific expression (diffASE)

Differential allele-specific expression (diffASE) was computed by comparing divergence in allele counts across control and burrowing conditions via a Fisher’s exact test. Since five of the six animals studied were related (3 siblings: 2 control, 1 burrowing; 2 siblings: 1 control, 1 burrowing; 1 outlier: burrowing), we were able to partially factor in genetic background by performing this analysis on two sets of siblings from opposite conditions and a third unrelated pair. For each pair, we compared allelic counts in a 2×2 contingency table (*P. polionotus* burrowing allele, *P. polionotus* control allele; *P. maniculatus* burrowing allele, *P. maniculatus* control allele). To assess significance across all three pairs, we combined the resulting *p-*values using Fisher’s method and then adjusted for multiple tests using Bonferroni correction. We also generated a more conservative gene list by filtering for situations in which all three replicates displayed significant diffASE (as opposed to significance from the combined *p-*values).

Since RNA-seq data can be prone to overdispersion, we also calculated diffASE using a beta-binomial model via the R package MBASED^29^. Analyses were performed using a 2-sample analysis with default parameters. Significance was assessed via *p*-values extracted from running 1,000,000 simulations per pair of animals (following the same design as the Fisher’s exact tests above). We found that, for each pair, the *p*-values calculated by MBASED were correlated with the Fisher’s exact test results at an r^2^>0.99, suggesting that both methods were capturing the same statistical structure in the data.

Using the genes identified as possessing significant diffASE (above), we performed analyses of allelic induction. We inferred the direction of allelic induction by comparing the log_2_ ratios of allelic counts as a function of *condition*, as opposed to species, such that, for each gene, two ratios were calculated: (1) log_2_(*P. maniculatus* burrowing/ *P. maniculatus* control) and (2) log_2_(*P. polionotus* burrowing/ *P. polionotus* control). We then categorized genes as having either *P. maniculatus-* or *P. polionotus-*biased induction by comparing difference in the absolute magnitude of these two ratios. To perform sign tests, we used the resulting lists as input for gene set-specific enrichments of either *P. maniculatus* or *P. polionotus* induction. To do so, we used a Fisher’s exact test to compare the number of upregulated *P. maniculatus* alleles and the number of upregulated *P. polionotus* alleles contained within a gene set. Using Bonferroni correction, we adjusted all *p-*values to control for multiple testing across gene sets in an ontology.

### Intersecting QTL and gene expression data

To test for possible intersection between *cis*-regulatory divergence and interspecific genetic variation we compared gene expression patterns to a published QTL map of burrowing from backcross of *P. polionotus* and *P. maniculatus* (Weber et al., 2013). The original QTL study associated entrance tunnel length, burrow length, and escape tunnel presence with 526 genetic markers, measuring the strength of association via log of odds (LOD). Here, we considered an expanded set of structural traits related to burrowing (n = 23) measured from the original data set. We explored the extent to which genes involved in locomotor coordination (identified by the burrowing-induced allele-specific induction analyses) possessed systematically different LOD scores compared to genomic background. To do so, we associated each gene from our set of interest with its nearest genomic marker (choosing just one gene in cases in which multiple genes were associated with the same maker) and, for each burrowing trait, computed the median LOD score for the gene set. We then computed the median and mean LOD scores for 10,000 randomly chosen sets of markers of the same size as the gene set of interest and calculated a *p-*value by calculating the number of times the shuffled LOD scores were greater than the observed one, divided by the number of permutations (n = 10,000). This test was repeated for seven different gene lists: differentially induced locomotor coordination genes (*P. polionotus-* and *P. maniculatus-*biased genes combined; n = 31 genes), *P. polionotus-*biased locomotor coordination genes (n = 26), *P. maniculatus-*biased locomotor coordination genes (n = 5), differentially induced genes associated with abnormal gait (n =22), genes with significant diffASE (n = 177), genes with the most significant ASE across all three burrowing replicates (measured via binomial test of allele counts; n = 200), and genes most differentially expressed between burrowing *P. maniculatus* and *P. polionotus* parental samples (n = 200).

To assess the ability of markers associated with locomotor coordination to predict the burrowing phenotypes, we complemented the LOD analyses with a series of multiple regression-based permutation tests. Here, we again used permutation tests (n = 10,000) on the 23 burrowing traits but instead compared the fit of a multiple regression predicting phenotypic measurements from the genotype at each gene list-associated marker for all 272 animals in the data set. We first computed the empirical fit (r^2^) for the markers associated with the gene set of interest and then extracted the fits for 10,000 random subsets of markers of the same length from which *p-*values were calculated above. Results for the tests described in this section are in Table S2-3.

### Quantification and Statistical Analysis

Statistical details of experiments can be found in results and corresponding methods details. All statistical tests were performed in R Version 3.6.1 (R Core Team, 2015).

## Author Contributions

R.A.Y., H.B.F., and H.E.H. conceived the study. C.K.H, H.C.M., and N.L.B collected and analyzed behavioral data. C.K.H., H.C.M., and N.L.B prepared samples for RNA sequencing. R.A.Y. processed and performed RNA-seq and QTL-intersection analyses. R.A.Y, C.K.H, H.B.F. and H.E.H. performed additional analysis and prepared the manuscript.

## Acknowledgements

We thank Carlo Artieri, Rachel Agoglia, Caitlin Lewarch and members of the Fraser and Hoekstra labs for helpful discussion and input. Caitlin Lewarch assisted with dissections; Kyle Turner assisted RNA-seq library preparation; Nimrod Rubinstein provided imprinted gene resources; and Nina Sokolov helped with behavior assays. This work was supported by a Stanford Bio-X, Stanford CEHG, and Stanford School of Medicine Deans Office to R.A.Y.; a Chapman Memorial Scholarship, a National Science Foundation (NSF) Graduate Research Fellowship, a Doctoral Dissertation Improvement Grant (NSF 1209753), and an American Fellowship from the American Association of University Women to H.C.M.; a Natural Sciences and Engineering Research Council of Canada Graduate Fellowship to N.L.B.; and a Beckman Young Investigator Award to H.E.H. H.E.H. is an Investigator of the Howard Hughes Medical Institute.

## Supplemental information

**Figure S1.**
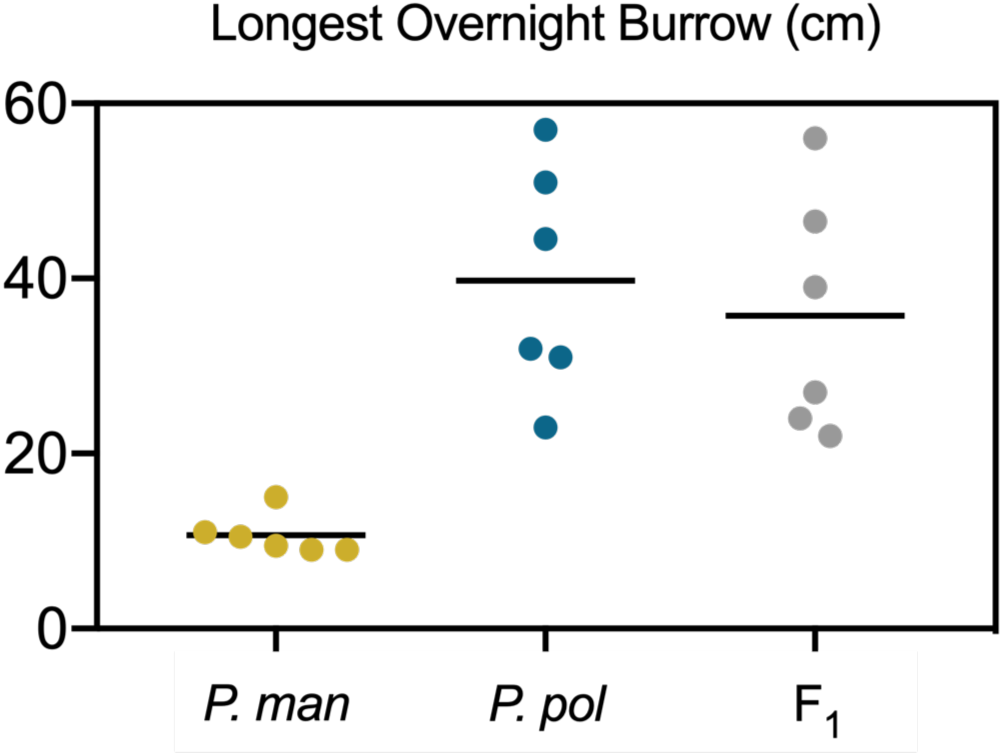
Longest overnight burrow of experimental animals. Each point represents the longest burrow (cm) of three trials dug by an individual of *P. maniculatus* (N=6), *P. polionotus* (N=6) or an F1 hybrid (N=6). Bar represents the mean burrow length.

**Figure S2.**
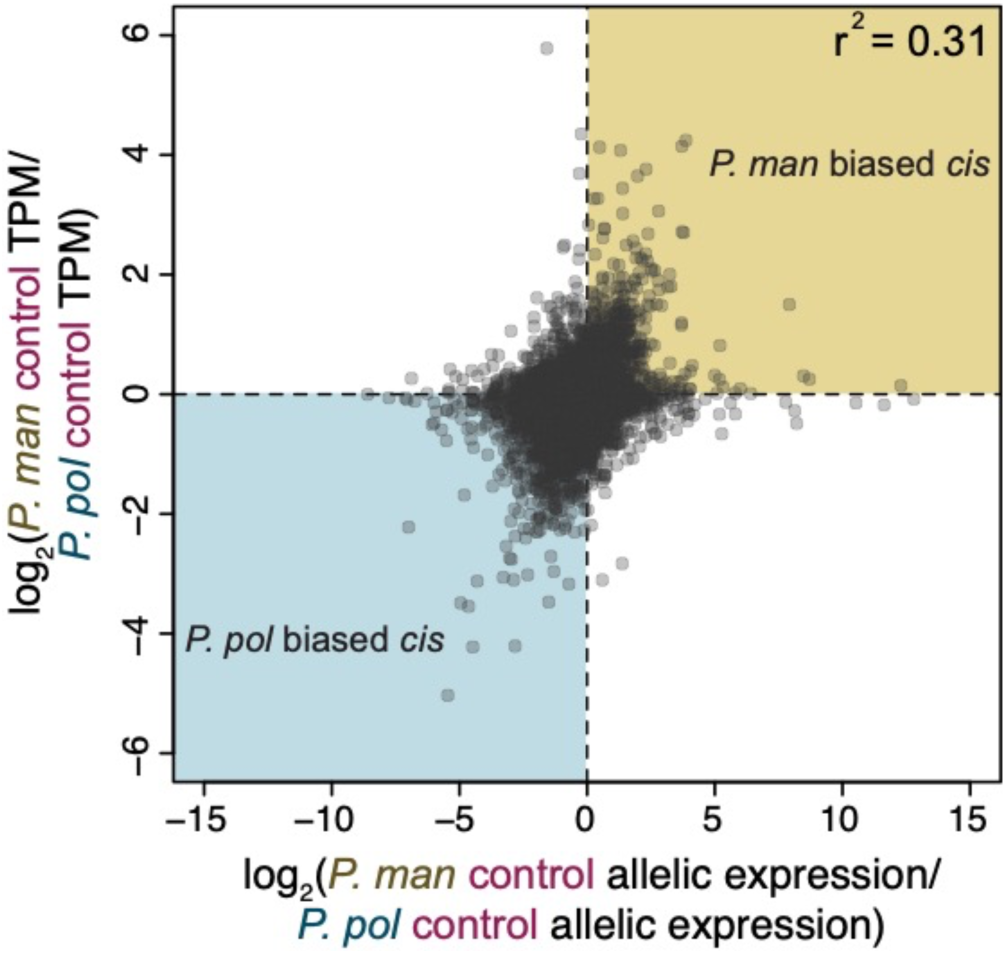
Comparison of the distribution of F_1_ hybrid allelic ratios in control animals (log_2_(*P. maniculatus* allele*/ P. polionotus* allele)) to the ratios of pure species control expression (log_2_(*P. maniculatus* TPM*/ P. polionotus* TPM)).

**Table S1.**
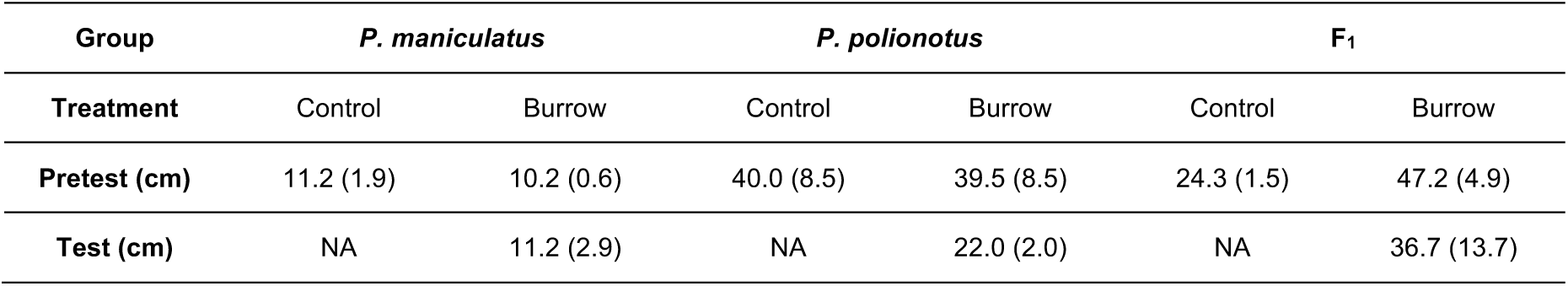
Burrowing behavior of experimental animals. Mean (SE) of the longest excavations dug by each species and F_1_ hybrids is shown.

**Table S2.** Results of the LOD based gene set comparisons. The LOD scores for markers associated with each gene set (labeled in the column headers) were compared to the distributions of LOD scores from 10,000 random permutations of markers for each of the three traits (row names). The *p-*values above were calculated by comparing the observed mean LOD score for the gene set of interest to the permuted distribution. ^*^ *p*<0.05, ^**^*p*<0.01, ^***^*p*<0.001, ^****^*p*<0.0001.

| Trait | Locomotor coordination (PO) | Locomotor coordination (BW) | Abnormal gait | Locomotor coordination (All) | diffASE | ASE | Parental differential expression |
| --- | --- | --- | --- | --- | --- | --- | --- |
| Total length | 0.0255<br>(*) | 0.0193<br>(*) | 0.0089<br>(**) | 0.0038<br>(**) | 0.0381<br>(*) | 0.1578 | 0.5101 |
| Average entrance tunnel length | 0.0136<br>(*) | 0.0119<br>(*) | 0.0167<br>(*) | 0.0004<br>(***) | 0.2346 | 0.6917 | 0.1552 |
| Average escape tunnel length | 0.4046 | 0.6384 | 0.4127 | 0.4143 | 0.5932 | 0.7021 | 0.9972 |

**Table S3.** Results of the regression-based gene set comparisons. The genotypes at markers associated with each gene set (labeled in the column headers) were used to predict measurements for the three traits via regression. The observed fits of these models were then compared to fits of regressions from 10,000 random permutations of markers. ^*^ *p*<0.05, ^**^*p*<0.01, ^***^*p*<0.001, ^****^*p*<0.0001.

| Trait | Locomotor<br>coordination<br>(PO) | Locomotor<br>coordination<br>(BW) | Locomotor<br>coordination<br>(All) |
| --- | --- | --- | --- |
| Total<br>length | 0.4791 | 0.4883 | 0.2022 |
| Average entrance<br>tunnel length | 0.4537 | 0.4597 | 0.3054 |
| Average escape<br>tunnel length | 0.00009<br>(***) | 0.1259 | 0.0009<br>(***) |

**Table S4.**
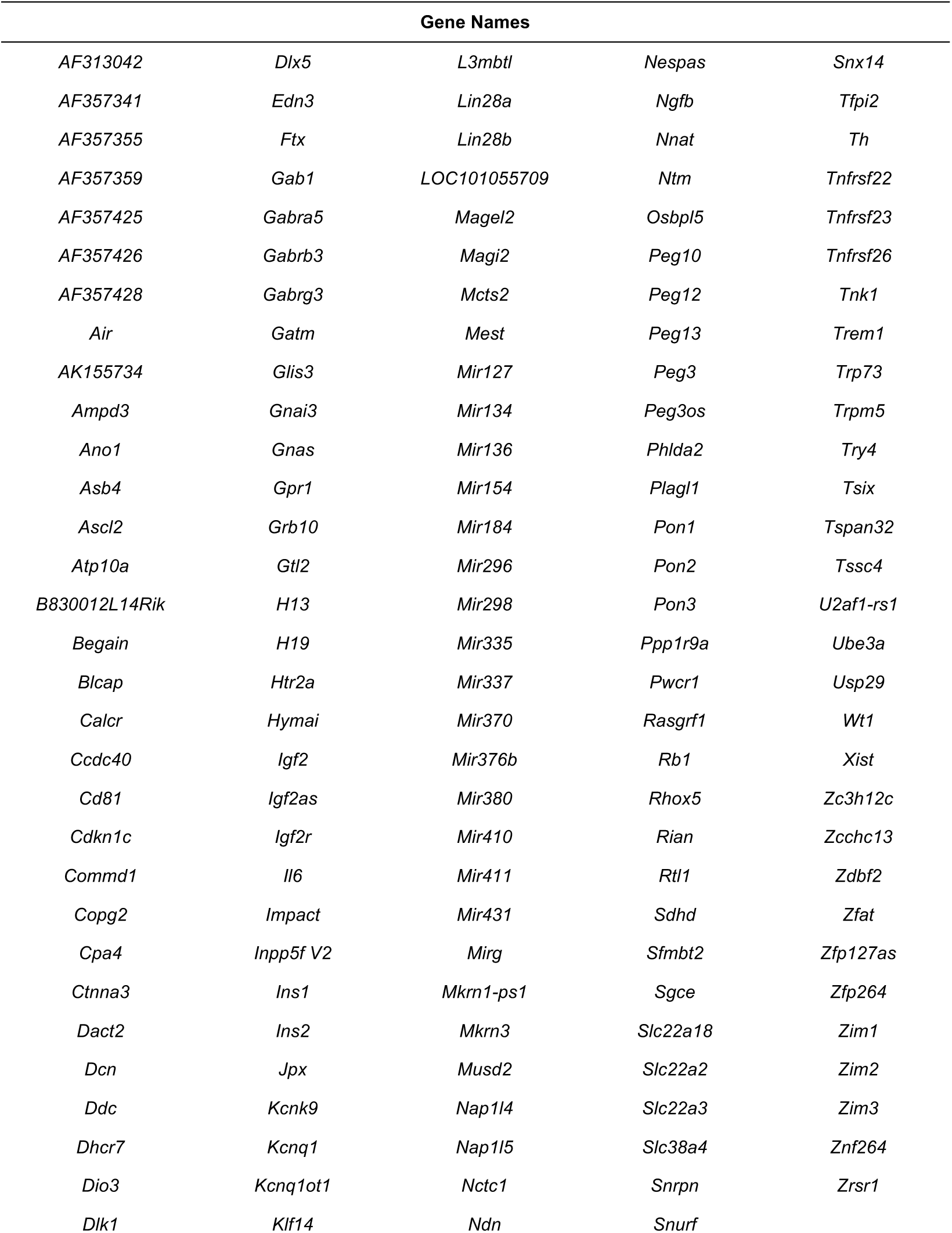
Alphabetized list of imprinted genes (N=154) removed from allele-specific expression analysis.

